# FPfilter: A false-positive-specific filter for whole-genome sequencing variant calling from GATK

**DOI:** 10.1101/2020.03.23.003525

**Authors:** Yuxiang Tan, Yu Zhang, Hengwen Yang, Zhinan Yin

## Abstract

1.

**Motivation:** As whole genome sequencing (WGS) is becoming cost-effective progressivelly, it has been applied increasingly in medical and scientific fields. Although the traditional variant-calling pipeline (BWA+GATK) has very high accuracy, false positives (FPs) are still an unavoidable problem that might lead to unfavorable outcomes, especially in clinical applications. As a result, filtering out FPs is recommended after variant calling. However, loss of true positives (TPs) is inevitable in FP-filtering methods, such as GATK hard filtering (GATK-HF). Therefore, to minimize the loss of TPs and maximize the filtration of FPs, a comprehensive understanding of the features of TPs and FPs, and building an improved model of classification are necessary. To obtain information about TPs and FPs, we used Platinum Genome (PT) as the mutation reference and its 300× deep sequenced dataset NA12878 as the simulation template. Then random sampling across depth gradients from NA12878 was performed to study the depth effect.

**Results:** FPs among heterozygous mutations were found to have pattern distinct from that of FPs among homozygous mutations. FPfilter makes use of this model to filter out FPs specifically. We evaluated FPfilter on a training dataset with depth gradients from NA12878 and a test dataset from NA12877 and NA24385. Compared with GATK-HF, FPfilter showed a significantly higher FP/TP filtration ratio and F-measure score. Our results indicate that FPfilter provides an improved model for distinguishing FPs from TPs and filters FPs with high specificity.

**Availability:** FPfilter is freely available for download on GitHub (https://github.com/yuxiangtan/FPfilter). Users can easily install it from anaconda.

## 2. Introduction

Whole-genome sequencing (WGS) is a technology that involves high-throughput sequencing platforms to sequence the whole genome DNA in individuals [1]. Compared with whole exon sequencing, WGS covers the whole genome (including exons, introns and intergenic regions) thus providing broader information for screening of pathogenic and susceptibility-associated gene loci and even for inferences about population evolution [2]. With the increase in sequencing throughput and further reductions in costs, WGS has been increasingly applied in health care and scientific research.

Currently, the most widely accepted WGS pipeline includes BWA-MEM for sequence alignment [3] and the variant-calling tool GATK [4]. However, recent studies showed that although the accuracy of this pipeline is already high, false positives (FPs) are still unavoidable [5]. As a result, after variant calling, filtering out FPs is recommended. Generally, filtering methods are subdivided into hard filtering, such as GATK-HF, and a machine learning-based method, such as GATK Variant Quality Score Recalibration (VQSR). GATK-HF can process single samples by setting fixed filtering thresholds of parameters and can run faster, whereas GATK VQSR requires at least 30 samples to build a model and takes more time.

Theoretically, it is impossible to identify true positives (TPs) and FPs in actual samples. Nonetheless, with the advent of reference samples from high-confidence reference mutagenesis datasets [Genome in a Botttle (GIAB) [6, 7] and Platinum genotypes(PT) [8]], it is now possible to perform TP and FP evaluation for algorithms. In the present study, PT was selected as the high-confidence reference dataset, because it has higher coverage and more mutations than GIAB does. To study the depth effect on FP features, we performed random sampling across gradients (from 17.8× to 71.2×) from the 300× deep-sequenced reference sample NA12878 and analyzed TPs and FPs after filtering.

Here, we present FPfilter, a novel tool for hard filtering of FPs with high specificity for GATK results from WGS data. By classifying variatnts into groups (SNPs v.s. INDELs and homozygous v.s. heterzygous) with distinct filters, FPfilter showed a significantly higher filtration ratio (the number of FP filtered to the number of TP filtered) and F-measure than GATK-HF. While filtering the same number of FPs, FPfilter can lose fewer TPs than GATK-HF can. This finding was proved on both the training dataset from NA12878 and on the testing sample from NA12877 and NA24385. Thus, FPfilter offeres an improved method to filter out FPs with fewer TPs lost.

## 3. Methods

### 3.1 Data simulation for performance evaluation

A deep-sequenced (300×) female WGS dataset (NA12878: SRR2052337 - SRR2052423) served as the simulation template. It was sequenced on the HiSeq2500 platform from the National Institute of Standards and Technology (NIST) [7].

Random subsamples of lower depths (17.8×, 35.6×, 53.4×, 71.2×,) were generated by the seqtk[9] tool with 3 replicates at each depth.

### 3.2 Reference genome and high-confidence mutation reference

GRCh37 from NCBI served as the reference genome.

After comparison with GIAB[5], PT[8] was chosen as the “gold standard” mutation reference, because PT has higher coverage and more mutations than GIAB does (see Supplemental Table 1).

### 3.3 Analysis pipeline

#### 3.3.1 Alignment to the reference genome

The bwa mem[3] served as the aligner, with multi-node acceleration using the GLAD system. Samtools[10] software help us to sort the bam files, and sambamba[11] to merge the bam files and index data. Picardtools[12] was employed to remove the PCR duplicates in the bam files. Finally Qualimap2[13] software was used for statistical analysis of mapping.

#### 3.3.2 Variant calling

We chose the GATK Haplotypecaller[14] to call mutations with a default pre-filter of QUAL> 30.

#### 3.3.3 Statistical evaluation by means of the PT reference mutation

The performance of algorithms was evaluated by means of the PT reference in software rtgtools[15] and vcfeval [16].

The F-measure was calculated from TPs, FPs and false negatives (FNs).

F-measure = 2TPs / (2TPs + FPs + FNs)

All software and database versions used in this study and their uses are summarized in the Supplemental Table 2.

### 3.4 Filtering procedure

Nine features (ADR, DP, FS, GQ, MQRankSum, QUAL, QD, ReadPosRankSum and SOR, see Supplemental Table 3 for description) were employed in both filters.

#### 3.4.1 FPfilter protocol

After classification into four types [homozygous single-nucleotide polymorphisms (SNPs), homozygous insertion-deletions (INDELs), heterozygous SNPs and heterozygous INDELs], mutations that met the following conditions were filtered as FP (see Supplemental Figure 1 for the workflow):

heterozygous INDELs: ADR > 3 and MQRankSum ≤ −1; MQRankSum > 3 and QD > 25; DP > 2000

heterozygous SNPs: ADR > 5 and MQRankSum < −2; ADR < 0.5 and MQRankSum > 3; ADR < 0.2 and MQRankSum > 0.5; MQRankSum > 5 and QD > 10; MQRankSum > 0 and QD > 30; MQRankSum < −9.5 and QD > 20; QUAL > 4000 homozygous INDELs: GQ < 12

homozygous SNPs: GQ < 6

#### 3.4.2 GATK-HF protocols

Mutations that met the following conditions were filtered out as FPs: INDEL: QD < 2.0; FS > 200.0; ReadPosRankSum < −20.0; SOR > 10.0 SNP:QD < 2.0; FS > 60.0; MQ < 40.0; SOR > 3.0; MQRankSum < −12.5; ReadPosRankSum < −8.0

### 3.5 Computational resources

A Genes’ Mind 500 server with 5 nodes, 192 GB memory, 80 TB HDD, 2TB SSD hard disk and 70 CPU cores was employed in this study.

## 4. Results

### 4.1 Homozygous and heterozygous mutations have different FP signatures

As shown in Supplemental Figure 2A, heterozygous FPs had higher ADRs than TPs did among both SNPs and INDELs, but homozygous FPs did not. As shown in Supplemental Figure 2B, most heterozygous FPs were at the tail of the Depth-QUAL distribution, while FPs and TPs were hard to separate by this feature combination among homozygous mutations. As a result, to filter FPs specifically by means of these features, we treated heterozygous and homozygous mutations separately in FPfilter.

### 4.2 FPfilter filters out FP calls with high specificity

FPfilter showed a higher FP/TP filtration ratio than GATK-HF did among both SNPs and INDELs (Figure 1). Among SNPs (Figure 1A), GATK-HF filtered out more than 10-fold more TPs than FPs at 17.8× depth. The fold difference in fitration between TPs and FPs got smaller as sequencing depth increased. Nonetheless, at 71.2× depth, the number of FPs filtered by GATK-HF constitured only 18.98% of the number of TPs filtered. On the contrary, FPfilter filtered out more FPs than TPs. Its FP/TP filtration ratio ranged from 2.59 to 6.09. The peak of the ratio was at 35.6× depth and the ratio slightly decreased as sequencing depth increased, indicating that the number of TPs filtered out increased as the sequencing depth increased. Among INDELs (Figure 1B), the FP/TP filtration ratio of FPfilter was still significantly higher than that of GATK-HF, but with less difference than that seen among SNPs. The FP/TP filtration ratio inceased as the sequencing depth increased, implying that higher sequencing depth kept benefiting INDEL calling.

**Figure 1:**
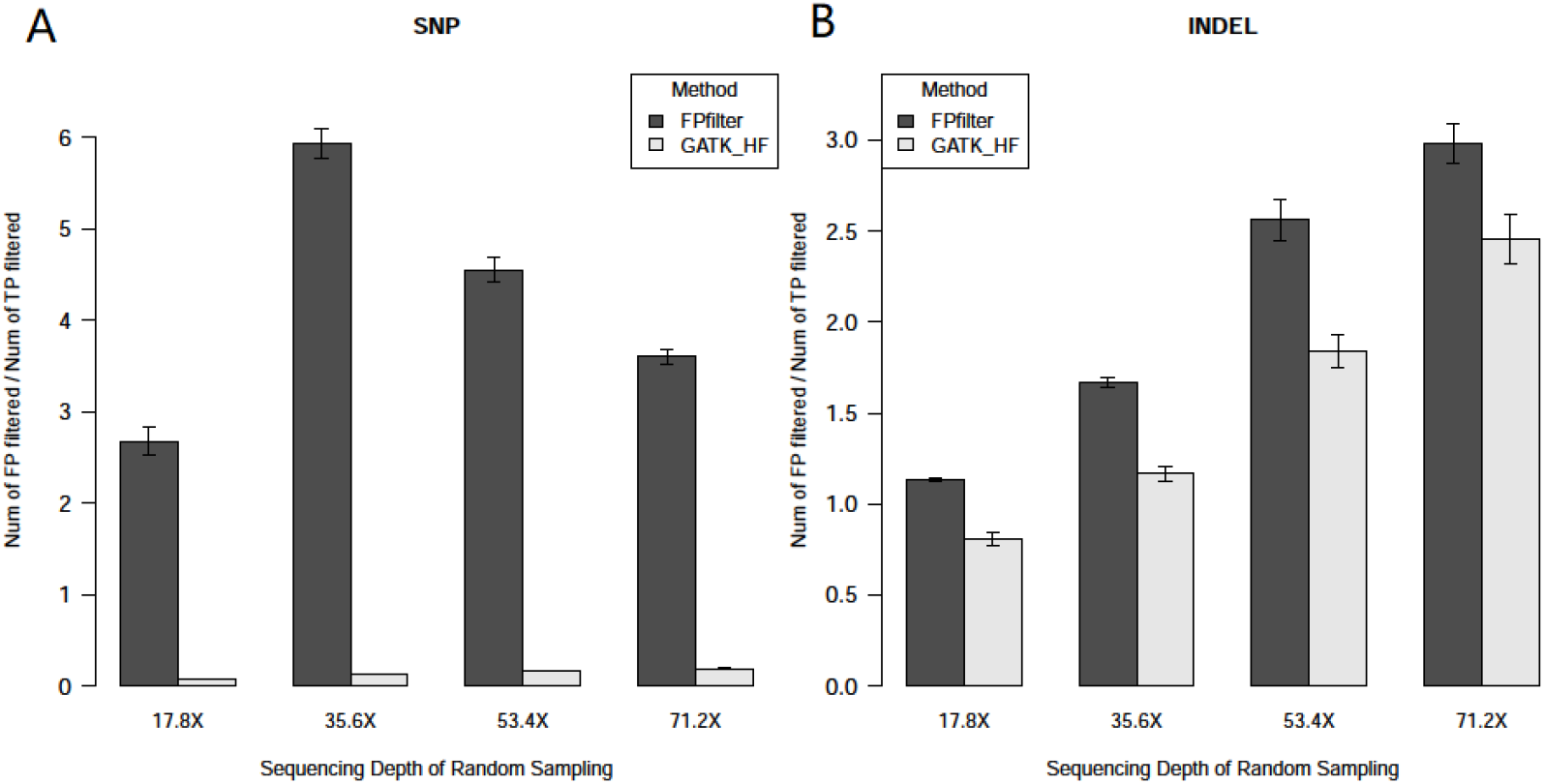
FPfilter has a higher ratio of FPs filtered to TPs filtered than GATK-HF does. Panel A presents SNP data; panel B presents INDEL data. The Y-axis represents the FP/TP filtration ratio (the number of FPs filtered divided by the number of TPs filtered). Black bars stand for the results from FPfilter; grey bars denote the results from GATK-HF. Error bars represent the standard error.

### 4.3 FPfilter shows a higher F-measure score than GATK-HF does

Although the absolute value of the difference was small (<0.01), FPfilter yield a significantly higher F-measure score than GATK-HF did except at 17.8× depth among INDELs (Figure 2 and Supplemental Table 4). The overall performance seems to be satuated among SNPs at ~53.4× depth, but not among INDELs. Both FPfilter and GATK-HF showed better performance on filtering SNPs than INDELs.

**Figure 2:**
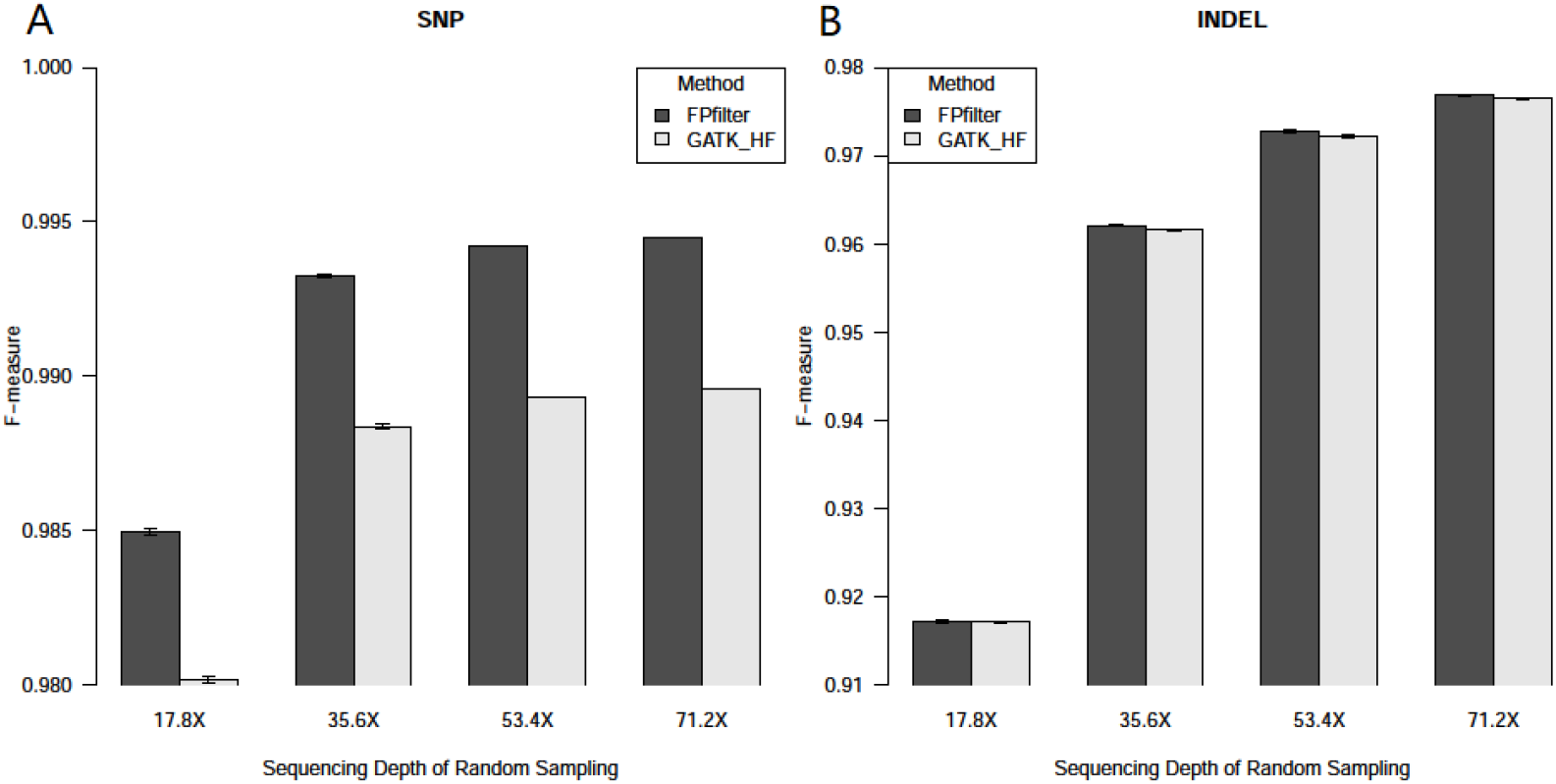
FPfilter yields a higher F-measure score than GATK-HF does. Figure A is on SNP; Figure B is on INDEL. Y-axis is the F-measure score. Black bar stands for result from FPfilter; grey bar stands for result from GATK-HF. Error bars represent the standard error.

### 4.4 FPfilter shows consistently better performance on a test sample than GATK-HF does

The ERR194146 sample from NA12877 was chosen as the test dataset. Although the FP/TP filtration ratio was smaller than that in the samples from NA12878, FPfilter outperformed GATK-HF on both the FP/TP filtration ratio and F-measure.

**Table 1.** Performance of GATK-HF and FPfilter on the ERR194146 sample

| Project |  | GATK-HF |  | FPrfilter |  |
| --- | --- | --- | --- | --- | --- |
| ID | Type | The FP/TP | F-measure | The FP/TP | F-measure |
| (Depth) |  | filtration ratio |  | filtration ratio |  |
| NA12877 | SNPs | 0.0332 | 0.9851 | 1.0875 | 0.9942 |
| (~50x) | INDELs | 0.2075 | 0.9731 | 0.7238 | 0.9759 |
| NA24385 | SNPs | 1.158422 | 0.9915 | 41.83721 | 0.9922 |
| (~60x) | INDELs | 0.402386 | 0.9307 | 0.909328 | 0.9313 |
| NA24143 | SNPs | 0.166109 | 0.9944 | 11.60131 | 0.9972 |
| (~60x) | INDELs | 0.172524 | 0.9779 | 1.419355 | 0.9795 |
| NA24631 | SNPs | 0.0739 | 0.9659 | 6.3262 | 0.9799 |
| (~80x) | INDELs | 0.1153 | 0.9593 | 0.5791 | 0.9684 |
| NA24695 | SNPs | 0.166109 | 0.9944 | 11.60131 | 0.9972 |
| (~60x) | INDELs | 0.172524 | 0.9779 | 1.419355 | 0.9795 |

可以考虑把高低深度的结果也加进来？

## 5. Conclusion & Discussion

Using a high-confidence mutation reference set (such as PT) and random subsamples across depth gradients (from 17.8× to 71.2×) of the 300× deep-sequenced actual data from NA12878, we systematically evaluated the features of TP and FP data among mutations, and found a useful feature (ADR) and an FP clustering model among heterozygous mutations (See Supplemental Figure 3). Besides that, we found that homozygous and heterozygous mutations have distinct signatures, which were never reported before.

Based on these findings, we developed an FP filtering tool: FPfilter. FPfilter filtered out FP mutations specifically, while losing fewer TPs and showing a higher F-measure score than GATK-HF did, on both the training dataset from NA12878 and the testing sample from NA12877. FPfilter ran ~1/3 faster than GATK-HF did with these samples, potentially because GATK-HF requires reference genome information in the process. However, because the total run time was less than 10 minutes per sample for both methods, we assumed that the run time of the filtering step is not a limiting factor in the WGS analysis procedure.

Although in the regular sequencing depth range of WGS (from 17.8× to 71.2×), the improvement on the F-measure by FPfilter was less than 1% as compared with that for GATK-HF, this improvement was meaningful, because the F-measure of GATK-HF was greater than 98% already. Meanwhile, FPfilter and GATK-HF were designed for analyzing stand-alone samples, such as those in personalized medicine. Any small improvement on finding TPs and/or filtering FPs might save a patient.

Furthermore, the original sum of mutation calls (including TPs, FPs and FNs) were around 4,000,000 depending on the sequencing depth. As a result, 1% of the total was not a small number. In conclusion, this small improvement by FPfilter could be clinically significant.

However, because of the limited number of high-confidence references, the evaluation of FPfilter on the test sample had weak in statistical power. Further demonstration might require a systematic experimental evaluation on real-world samples, which was not affordable in this study.

Besides GATK-HF, we tested GATK-VQSR on both training and testing datasets, but failed to find enough samples to build the Gaussian mixture model. Because the main use of FPfilter is single-sample processing, we only compared it with GATK-HF in the end.

It is worth noting that, the filtration criteria of FPfilter and GATK-HF hardly overlap. If a user wants to filter out FPs exhaustively without consideration of the loss of TPs, he/she can use FPfilter together with GATK-HF. Moreover, compared with GATK-HF, FPfilter is much less aggressive on filtering FPs in SNPs but more aggressive on filtering FPs in INDELs (Supplemental Table 5). Therefore, to have appropriate filtering, users can utilize only one of the filtered results of SNPs or INDELs. 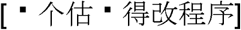 On the opposite, if a user focuses only on the number of TPs, no post-call filtering should be done.

We also noticed that other variant calling methods, such as VariantMetaCaller[17], freebayes[18] and DeepVariant[19], requiring little post-call filtering, were stated to perform better than GATK. But before their replacement of GATK, imporving the post-call filtering of GATK is still meaningful.

## 6. Conflict of Interest

The authors declare no conflicts of interest.

## 7. Acknowledgement

This work was supported by the China Postdoctoral Science Foundation (grant No. 2017M622901 to Dr. Yuxiang Tan); the Major International Joint Research Program of China (grant No. 31420103901); and the “111 project” (grant No. B16021) to Dr. Zhinan Yin and Hengwen Yang.

