## Supplemental information for "FPfilter: A false-positive-specific filter for whole-genome sequencing variant calling from GATK"

**Supplemental Table 1: Statistics on the comparison between GIAB and PT**

| Name | GIAB-NISTv3.3.2 NA12878 | Illumina Platinum Genomes NA12878 |
| --- | --- | --- |
| All mutations | 3775119 | 4049512 |
| SNPs | 3258078 | 3524621 |
| Insertions | 242384 | 256344 |
| Deletions | 261643 | 261754 |
| Indels | 13014 | 6793 |
| Phased Genotypes | 99.5% (3756728/3775119) | 100.0% (4049512/4049512) |
| Total Het/Hom ratio | 1.54 (2288911/1486208) | 1.60 (2490061/1559451) |

**Supplemental Table 2: Software and databases used in this study**

| Name of software/database | version | application |
| --- | --- | --- |
| aspera-connect | 3.1.1.70545 | Sra download |
| sratoolkit | 2.8.1 | Sra uncompress |
| fastqc | v0.11.5 | Fastq qc |
| BBMap | 37.10 | Fastq process |
| fastx_toolkit | 0.0.14 | Fastq process |
| ReSeqtools | 0.23 | Fastq qc |
| GRCh37 | - | Reference genome |
| GIAB | 3.3.2 | High-confidence reference mutation set |
| Platinum Genomes | V2016-1.0 | High-confidence reference mutation set |
| glad | v2.1 | Multi-node parallel acceleration system |
| bwa | 0.7.13-r1126 | Aligner |
| samtools | 1.3.1 | Bam process |
| sambamba | v0.6.4 | Bam process |
| picardtools | 1.119 | Bam process |
| GATK Haplotypecaller | 3.7 | Variant caller |
| rtg-tools vcfeval | 3.7.1-eb13bbb | Accuracy evaluation |
| R | 3.2.5 | Visualization |
| ggplot2 | 2.2.1 | Visualization |
| scatterplot3D | - | Visualization |

**Supplemental Table 3: Technical definitions of features of sequencing quality**

| Feature | Definition |
| --- | --- |
| AD | Also known as the allele depth, this term means the unfiltered count of reads that support a given allele for an individual sample. The values in the field are ordered to match the order of alleles specified in the REF and ALT fields: REF, ALT1, ALT2 and so on if there are multiple ALT alleles. |
| ADR | The ratio of the read depths between REF and ALT (the number of reads supporting the base of the reference genome +1) divided by (the number of reads supporting mutation + 1) |
| ALT | Alternate non-reference base(s) |
| Depth (DP) | Total (unfiltered) depth of coverage |
| FisherStrand (FS) | A phred-scaled p-value according to Fisher’s Exact Test(Fisher, 1922) to detect strand bias (the variation being seen on only the forward or only the reverse strand) in the reads. More bias is indicative of false positive calls. |
| GQ | Conditional genotype quality, encoded as a phred quality −10log10p |
| MQRankSum (MQRS) | The u-based z-approximation from the Mann-Whitney rank-sum test(Mann and Whitney, 1947) for mapping qualities (reads with ref bases vs. those with the alternate allele). Note that the mapping quality rank sum test can not be performed on sites without a mixture of reads showing both the reference and alternate alleles. |
| QualByDepth (QD) | Variant confidence (from the QUAL field) divided by unfiltered depth of non-reference samples. |
| QUAL | Phred-scaled quality score for the assertion made in ALT |
| ReadPosRankSum | The u-based z-approximation from the Mann-Whitney rank-sum test(Mann and Whitney, 1947) for the distance from the end of the read for reads with the alternate allele. If the alternate allele is only seen near the ends of reads, this is indicative of an error. Note that the read position rank sum test can not be performed on sites without a mixture of reads showing both the reference and alternate alleles. |
| REF | Reference base(s) |
| RMSMappingQuality (MQ) | This is the Root Mean Square of the mapping quality of the reads across all samples. |
| StrandOddsRatio (SOR) | The StrandOddsRatio annotation is one of several methods are intended to test whether there is a strand bias in the data. Higher values indicate greater strand bias. |

**Supplemental Table 4 Statistical values of F-measure between two filters**

| Type | Depth | | FPfilter | | GATK-HF | |
| --- | --- | --- | --- | --- | --- | --- |
|  |  |  | Median | Standard Error | Median | Standard Error |
| SNP | | 17.8X | 0.984977473 | 8.92558E-05 | 0.980174377 | 7.04331E-05 |
|  |  | 35.6X | 0.993236741 | 0.988377353 | 2.25086E-05 | 3.25765E-05 |
|  |  | 53.4X | 0.994205387 | 9.37793E-06 | 0.989317892 | 8.99188E-06 |
|  |  | 71.2X | 0.994479378 | 5.48702E-06 | 0.989594686 | 5.32033E-06 |
| INDEL | | 17.8X | 0.917123404 | 0.00012819 | 0.917167405 | 0.000112121 |
|  |  | 35.6X | 0.962056571 | 6.50E-05 | 0.961528996 | 8.49E-05 |
|  |  | 53.4X | 0.972722067 | 0.000133736 | 0.972143688 | 0.000165439 |
|  |  | 71.2X | 0.976865145 | 5.13E-05 | 0.976463313 | 3.86E-05 |

**Supplemental Table 5 The median of lose on TP and FP between two filters**

| Type | Depth | FPfilter | | GATK-HF | |
| --- | --- | --- | --- | --- | --- |
|  |  | TP | FP | TP | FP |
| SNP | 17.8X | 596 | 1593 | 34268.5 | 2462.5 |
|  | 35.6X | 617 | 3648 | 35223 | 4560 |
|  | 53.4X | 1045 | 4738 | 36197 | 5943 |
|  | 71.2X | 1529 | 5498 | 36942 | 7012 |
| INDEL | 17.8X | 2740 | 3111 | 204.5 | 165.5 |
|  | 35.6X | 979 | 1608 | 412 | 490 |
|  | 53.4X | 580 | 1470 | 406 | 742 |
|  | 71.2X | 492 | 1482 | 390 | 925 |

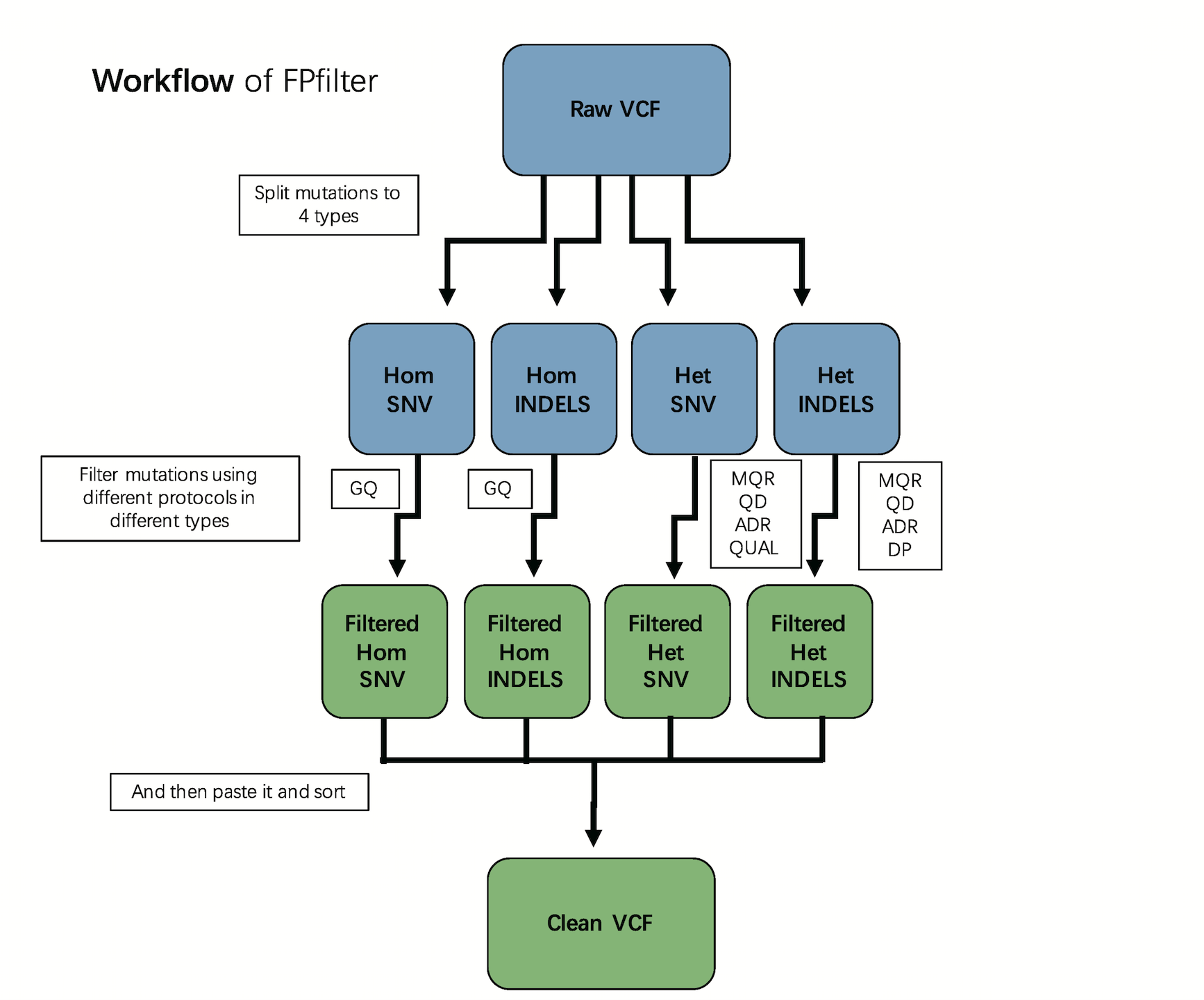

**Supplemental Figure 1: FPfilter workflow.**

First, the mutations are divided into four types; second, for each type, mutations are filtered by type-specific rules; finally, mutations after filtering are merged into one variant call format (VCF) file.

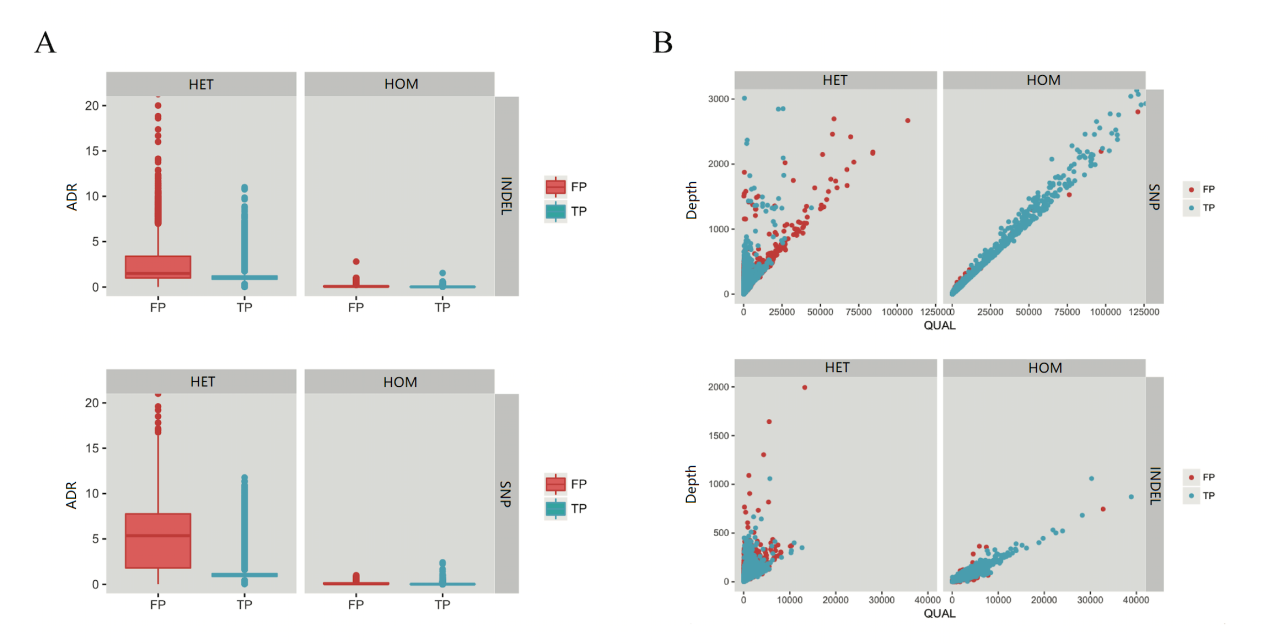

**Supplemental Figure 2: The two FP-specific models evaluated among heterozygous mutations at 71.2× depth as an example.**

A) In the ADR model, regardless of whether mutations are INDELs or SNPs, the ADRs were significantly higher among the FPs (red) than TPs (blue) among heterozygous (HET) mutations. However, among homozygous (HOM) locations, there is no significant difference between FPs and TPs. B) In the QUAL-Depth model, FPs can only be relatively separated among TPs in the heterozygous mutations consisting of SNPs.

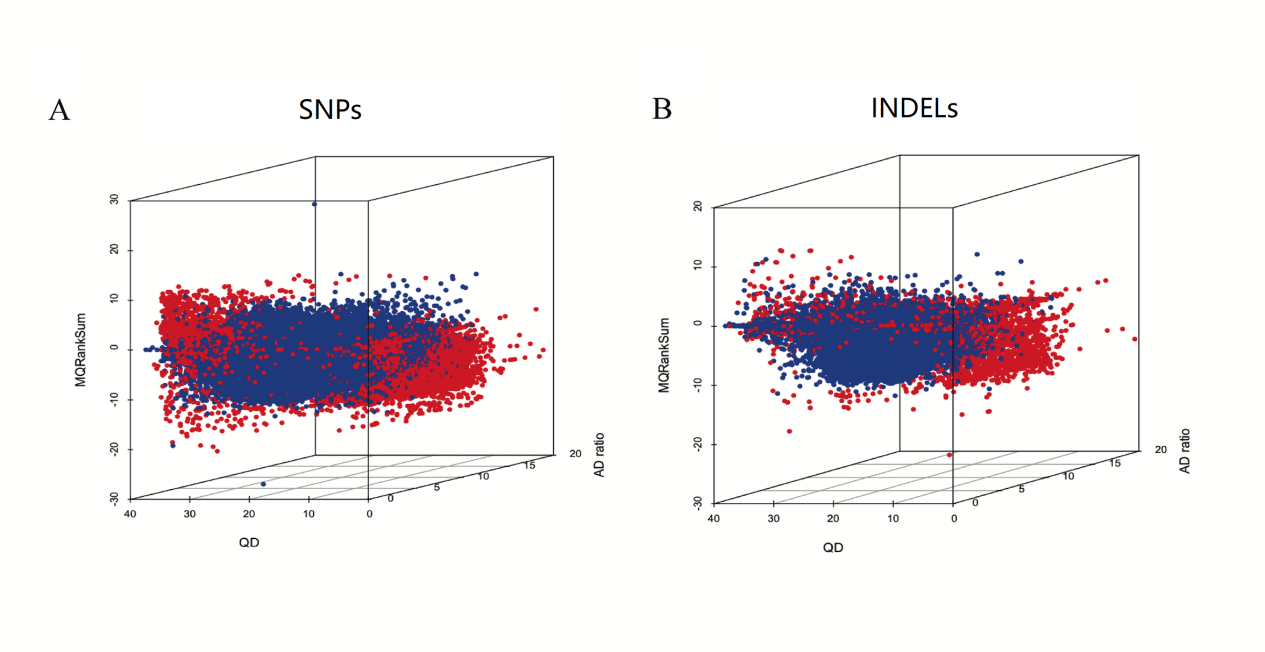
 **Supplemental Figure 3: QD-ADR-MQRS model of heterozygous locations at 71.2× depth as an example**.

A) The distribution of heterozygous SNPs; B) the distribution of heterozygous INDELs. The x-axis shows QD, the y-axis presents ADR, and the z-axis denotes MQRankSum (MQRS). Red dots represent FPs, and blue dots represent TPs.
